# Wolf-dog hybrids arise: new insight into the eastern fringe of the Northern European wolf population

**DOI:** 10.1101/2023.12.04.569906

**Authors:** Konstantin F. Tirronen, Anastasiia S. Kuznetsova

## Abstract

In this study, tissue samples of 35 wolves hunted in Russian Karelia and 12 specimens of dogs were used in 2012-2022 for genetic analysis. Four of the wolves (11%) were recognized by to phenotypic characteristics as wolf-dog hybrids (WDH). The analysis of 11 autosomal STRs confirmed the hybrid origin of three and excluded one. At the same time, two more admixed individuals with no phenotypic deviations were additionally identified, raising the total number of hybrids to five (14%). The analysis of CR1 mtDNA in the specified sample set revealed the dog haplotype in one of the phenotypically inferred WDH. The mtDNA haplotype diversity was represented by two haplotypes common in Eurasia. The genetic diversity level of the population revealed by 11 microsatellite loci was high (He = 0.746; Ho = 0.655; A = 8). However, the observed heterozygosity proved to be notably lower than expected, and the inbreeding coefficient was also high (Fis = 0.131) and, moreover, higher than the previously reported (Aspi et al. 2009). The wolf population of Karelia used to be considered as pure wolves, with a moderate level of hunting pressure. The observed genetic processes: interspecific hybridization and increased inbreeding occur against the background of a new wave of wolf fighting in Karelia. This region is of importance in the biogeographic sense – through this territory, the animal populations living in Scandinavia are connected to the Russian Plain, and further to Siberia.

## INTRODUCTION

In most cases, the animal’s phenotype is an important primary characteristic that a specialist or a layman can rely on in identifying the biological species. Everyone is well aware of the typical appearance of a wolf – a gray wolf; at least this characteristic is true for northern Eurasia. There are color differences between wolves of the Old and New Worlds, and black morphs are extremely rare in Eurasia as opposed to North America (Bibikov 1985; Yudin 2013; Apollonio et al. 2004; Pilot et al. 2019). It is claimed that the black color variation in wolves of America appeared as an introgression of dog’s genome in the past, and the genetic basis of inheriting of this sign has been studied quite closely but it still seems under discussion (Anderson et al. 2009; Hedrick 2009). We failed to find any mentions in the literature of black-colored animals in Fennoscandia. In February 2021, one black wolf (GW_244) was legally shot down in the south-west of Russian Karelia (RK), near Russian-Finnish border and a few km away from Karelia-Leningrad Region border (61°19’47” N 29°26’10” E). A curious fact personally communicated by Ilpo Kojola is that there is a transboundary pack in Finland at southern tip of the Finnish-Russian border in which many if not most individuals are black.

Interspecific hybridization between wolf and dog is a topical issue. Many studies across the world have examined various aspects and levels within this topic: local (Hindrikson et al., 2012; Molchan et al. 2023), worldwide or international (Pilot et al 2019), legal status (Trouwborst 2014; Salvatori et al. 2020), hybrid identification (Kusak et al. 2018; Dziech 2021), proposals on unifying genetic identification approaches (Randi et al. 2014; Lorenzini et al. 2022), and many others. The level of knowledge of the population genetic structure varies notably across the wolf range in Eurasia. Many European populations have been studied in sufficient detailed using genetic markers of various classes: mitochondrial DNA, autosomal and Y-linked microsatellites STR (Short Tandem Repeats), SNP (Single Nucleotide Polymorphism), and whole-genome analyses (Harmoinen et al. 2021; Hindrikson et al. 2016; Jansson et al. 2014; Pilot et al. 2007, 2018; Smeds et al. 2021; Sundqvis et al. 2001; Vilaça et al. 2023). At the same time, Russia’s wolf range has been poorly studied and wolf-dog hybridization is discussed in just a few studies (Korablev et al. 2021; Pilot et al. 2019).

The focus in this study is wolves of Russian Karelia (Republic of Karelia, RK). The importance of this population is in ensuring the connectivity between Fennoscandian wolves and the population of the East European Plain (Åkesson et al. 2021; Harmoinen et al. 2021). Studies of the neighboring Finland’s wolf population and samples from RK territory prove the need for further genetic study of the wolf population in this region (Aspi et al. 2009; Jansson et al. 2012). Having previously discussed the issues of wolf hunting in RK and investigating the diversity of mitochondrial haplotypes, we recorded a wild wolf-like animal with dog’s CR1 mtDNA haplotype (Tirronen et al. 2023). However, there have been no targeted studies of the genetic structure and wolf-dog hybridization in the RK population. Urged by this information and data on wolves with phenotype deviations, we decided to study possible cases of wolf-dog hybridization in RK.

We undertook to evaluate the consequences of the recent increase in hunting pressure for the wolf population, possibly manifested in interspecific hybridization, increased inbreeding, and changes in the genetic diversity and population structure. The tasks were to genetically assess whether there is evidence of mixing between wolves and dogs in Karelia using wild phenotypically deviated canids and a randomly assigned sample set.

## MATERIAL AND METHODS

### Study area and wolf population

All samples were collected in the time period of 2012-2022 from the territory of Russian Karelia, which is situated in north-western Russia and covered by boreal forest (Fig. 1). The study area spans for approximately 660 km north to south, and more than 420 km west to east. The total area of RK is 172400 km^2^ (excluding the White Sea area) and more than half of the territory is forested, almost a quarter is under waterbodies (rivers and lakes), a little less is under wetlands, and the rest is farmland, settlements, and industrial areas.

**Fig. 1.**
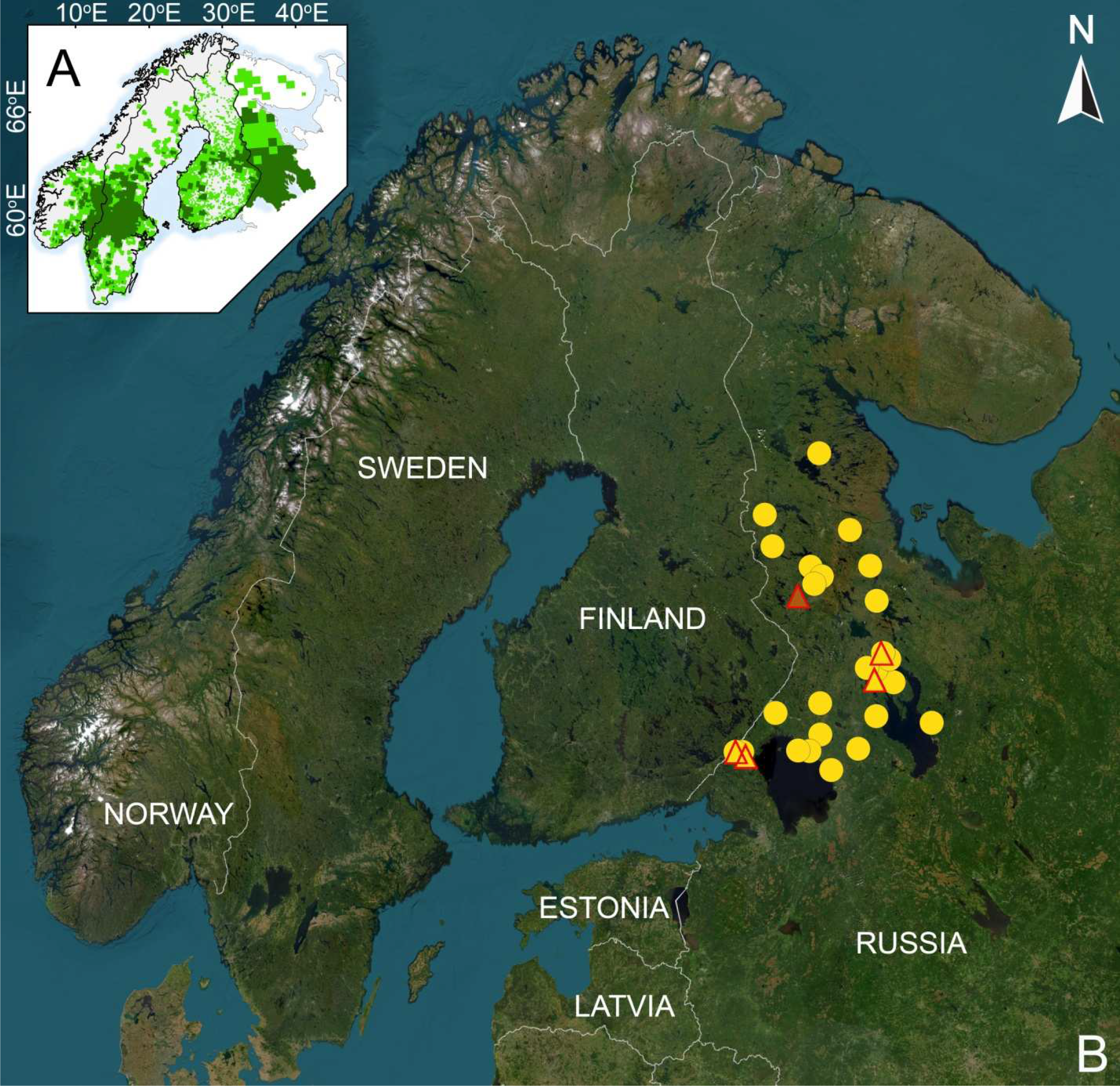
Wolves in Fennoscandia according to Kaczensky et al. (2021), supplemented by our data for the Murmansk and Karelia regions (A). Sampling locations in the Republic of Karelia (B): circles - pure wolves, triangles - wolf-dog hybrids (yellow color is the wolf CR1 mtDNA haplotype, brown - dog CR1 mtDNA haplotype)

A few remarks are due on the current status of the wolf in Karelia:

1. Population size. Since the beginning of the new millennium, the wolf population has ranged from about 280 to 450 individuals in February-March (the period of annual animal counts), i.e. at the end of the hunting season and before the offspring appear.
2. Distribution. Wolves live throughout RK, but the population is unevenly distributed – the population density in the north of the region (northern taiga subzone) is much lower than in the south (middle taiga subzone).
3. Wolf hunting. Since 2000, the annual harvest has varied from 45 to 236 animals. It did not exceed 100 before 2010, but since 2012 it has been steadily higher than 100, exceeded 160 in 2016 and reached an absolute peak of 236 individuals in 2018. In total, more than 2,700 wolves have been killed in Karelia since the beginning of the new millennium. A monetary bounty is paid to the hunter for each wolf they kill. Doubling of the bounties in 2016 increased the harvest. Since 2018, wolf hunters are additionally rewarded in the form of a license to hunt moose, which has significantly stimulated wolf hunting.
4. Phenotype. Since bounties are paid for wolf extermination, each hunted animal undergoes a veterinary examination including evaluation of correspondence between the animal’s body size and its approximate age and identification of morphological traits (color, head and tail shape, body proportions) corresponding to the biological species – the wolf.

### Sampling

In this study we used 47 canids samples, 36 of them belonging to wild and 11 to domestic animals. Among the 36 wild canid samples, 31 belong to wolves, four presumably to hybrids, and one to a feral dog. The set of wolf samples for genetic analysis, n = 35, was randomly selected (with the exception of four phenotypic hybrids), attempting to select samples that cover the territory of Karelia as uniformly as possible.

Before 2017, samples for genetic analysis accounted for 5-10% of the harvested individuals. Since then, the sampling rate was gradually raised to 85% of the annual wolf harvest in RK. With hunters supposed to receive bounties for each wolf they kill, where there uncertainty regarding animals with atypical “dog” traits, their samples were necessarily sent to our laboratory for additional genetic studies. Thus, samples of four suspected WDH were delivered to us.

All the wolves were legally killed and the samples were provided by the Hunting Department (RK Ministry of Natural Resources and Environment), which oversees hunting.

Muscle tissue or a piece of skin with muscle fibers were taken from the killed animals, placed in tubes with ethanol, and stored at −20°C until analysis.

To identify possible hybridization, 11 mongrel dogs were included in the analysis as a reference group. Samples from approximately “wolf-sized” dogs were obtained from a trapped dog facility for stray or homeless dogs who could, hypothetically, have participated in interspecific crossbreeding. Also, we used one sample of a feral dog (GW_50) trapped in the wild in the south-west of RK. Dog hairs were put in paper bags and stored in a dark and cold place until analysis.

All sampling locations, mtDNA haplotypes, genetic status of animals based on 11 autosomal microsatellites were entered in an Excel spreadsheet. Based on this data, a GIS of the study area incorporating the analyzed data was created using QGIS 3.4.14

### Microsatellite analysis

We used 20 microsatellite markers: CPH2, CPH4, CPH5, CPH8, CPH12 (Fredholm and Wintero 1995), FH2001, FH2017, FH2010, FH2054, FH2079, FH2088, FH2096 (Francisco et al., 1996), AHT130, AHT137 (Holmes et al., 1995), INRA21 (Mariat et al, 1996), AHTk211 (Lingaas et al, 1997), vWF (Shibuya et al, 1994), C09.173, CXX279, C20.253 (Ostrander et al, 1993). The forward primer for each locus was labeled with fluorescent dye: FAM, ROX, TAMRA, R6G. All loci were polymerase chain reaction (PCR) amplified in a volume of 25 μl containing 16.5 μL H2O, 5 μL Screen Mix (Evrogen), 1 μL of each primer (10 μM, Syntol), and 1.5 μL of matrix DNA. PCR protocol: initial denaturation at 95°C for 3 min, 35 cycles, including 20 s at 95°C, 20 s at 58-62°C, and 20 s at 72°C, and final elongation for 5 min at 72°C.

To identify the length of the amplified loci 0.5 μl of the molecular size standard GeneScanTM 600 LIZ (Applied Biosystems, USA) was added to PCR products.

The fragment length was determined using capillary electrophoresis on Seqstudio genetic analyzer (Applied Biosystems, USA) following the protocol provided by the manufacturer. The results were analyzed in GENEMAPPER v. 4.0 (Applied Biosystems, USA). Some samples were repeated twice to avoid genotyping errors. The presence of null alleles and stuttering were analyzed with MICRO-CHECKER v2.2. (van Oosterhout et al. 2004).

### CR1 mtDNA sequencing

Partial sequences (350 bp) of the hypervariable part of the mitochondrial DNA control-region (mtDNA CR1) (Saccone et al., 1987) were obtained using universal PCR primers Thr-L 15926 5’-CAATTCCCCGGTCTTGTAAACC-3’ and DL-H 16340 5’-CCTGAAGTAGGAACCAGATG-3’ (Vilà et al, 1999) from wild animals. The amplification of mtDNA control region I fragment was carried out the same way as in microsatellite analysis at 52°C annealing temperature. The nucleotide sequences of the amplified mtDNA region were determined according to the Sanger method in two directions using BigDye Terminator 3.1 DNA sequencing kits (Applied Biosystems, United States) on Seqstudio genetic analyzer (Applied Biosystems). The obtained sequences were edited manually and aligned in MEGA11 (Tamura et al, 2021) using a ClustalW algorithm. The haplotype (Hd) and nucleotide (Pi) diversity were calculated using the DnaSP 5.0 software (Librado and Rozas, 2009).

### Genetic diversity

GENALEX software (Peakall and Smouse 2012) was used to calculate the number of alleles per locus, observed heterozygosity (Ho), expected heterozygosity (He), and the test for the Hardy– Weinberg equilibrium (HWE) in wolves separately. The inbreeding coefficient (Fis) and its statistical significance were calculated using ARLEQUIN v.3.5 (Excoffier and Lischer 2010).

To detect the recent bottleneck signature in the Karelian wolf population, we used BOTTLENECK v1.2.02 software (Cornuet and Luikart 1996). The TPM mutation model was chosen as the most appropriate for microsatellite data and a probability of 95% for single-step mutations with a variance of 12 was applied as suggested by S. Piry et al. (1999). Since we had fewer than 20 loci, Wilcoxon’s test was used to check if significant excess of heterozygosity was detected (Luikart and Cornuet 1997). Also, a mode shift test was applied to check if the frequency classes based on allele size deviated from the normal L-shaped distribution, indicating rare allele loss.

To estimate relatedness or relationship between individuals on the basis of microsatellite data, we used the ML-RELATE software (Kalinowski et al. 2006). Simulations were performed in two types of hypothesis tests.

### Analysis of wolf-dog hybridization

To identify wolf-dog hybrids we used genotypes from both the wild wolf population and dogs. STRUCTURE v2.3.4 (Pritchard et al. 2000) was applied to evaluate the number of genetic clusters (K) in the data and to assign individuals to their likely origin. The dataset for identifying hybrid samples comprised all 47 individuals, including wolves (n = 31), dogs (n = 12), and four hybrids (n = 4). The run parameters were the following: “admixture” and independent allele frequency “I” models, without any prior population information, assuming K from 1 to 10. Five independent runs were done for each K using 1000000 sweeps of the Monte Carlo Markov chain (MCMC) and discarding the first 100000 as burn-ins. The results were then processed in STRUCTURE HARVESTER (Earl and vonHoldt 2012). The ΔK statistics were used to identify the highest rate of increase in the posterior probability LnP(D) of the data between each consecutive K as described by Evanno et al. (2005). CLUMPP (Jakobsson and Rosenberg 2007) was used to match the data from the multiple runs for each K and STRUCTURE Plot (Ramasamy et al. 2014) was used to display the results.

GENETIX v. 4.05 (Belkhir et al. 2004) was used to distinguish between wolves, dogs, and wolf-dog hybrids on the basis of microsatellite data with factorial correspondence analysis (FCA).

To test if these wolves were related, we calculated the maximum likelihood of relatedness using ML-RELATE program (Kalinowski et al. 2006).

NEWHYBRIDS v1.1 beta (Anderson and Thompson 2002) was used to calculate the posterior probability on MCMC estimates that the individual belongs to each of the 6 predefined ancestry classes: wolf, dog, hybrids F1 and F2, and respective first-generation backcrosses to wolf and dog. The overall data set was processed with 1000000 MCMC sweeps.

## RESULTS

### Phenotype

According to the phenotype descriptions of the entire set of samples, n = 35, four individuals were preliminarily assigned to the WDH category. The traits by which these individuals are distinguished from the wolf general set are unequal. GW_119 was a semi-synanthropic animal from the outskirts of a city with a population of about 14 thousand people. The distinctive traits were white claws and white pads. GW_244 was a completely black wolf-like animal (Fig. 2) that lived in a pack of typical wolves and was shot down together with an ordinary wolf of the same pack. Sample GW_246 was not supplied with an exact description, but was accompanied by a note “hybrid”. The adult female GW_494 had no black stripes on front legs, no black tip of tail, the overall color tone was brownish red, the shape of the head and ears resembled Akita Inu dog breed (Fig. 2). The sample set in which these four WDHs were found belongs to the period 2017-2022, during which 1127 wolves were harvested. Thus, the share of the presumed phenotypic hybrids is 0.35%.

**Fig. 2.**
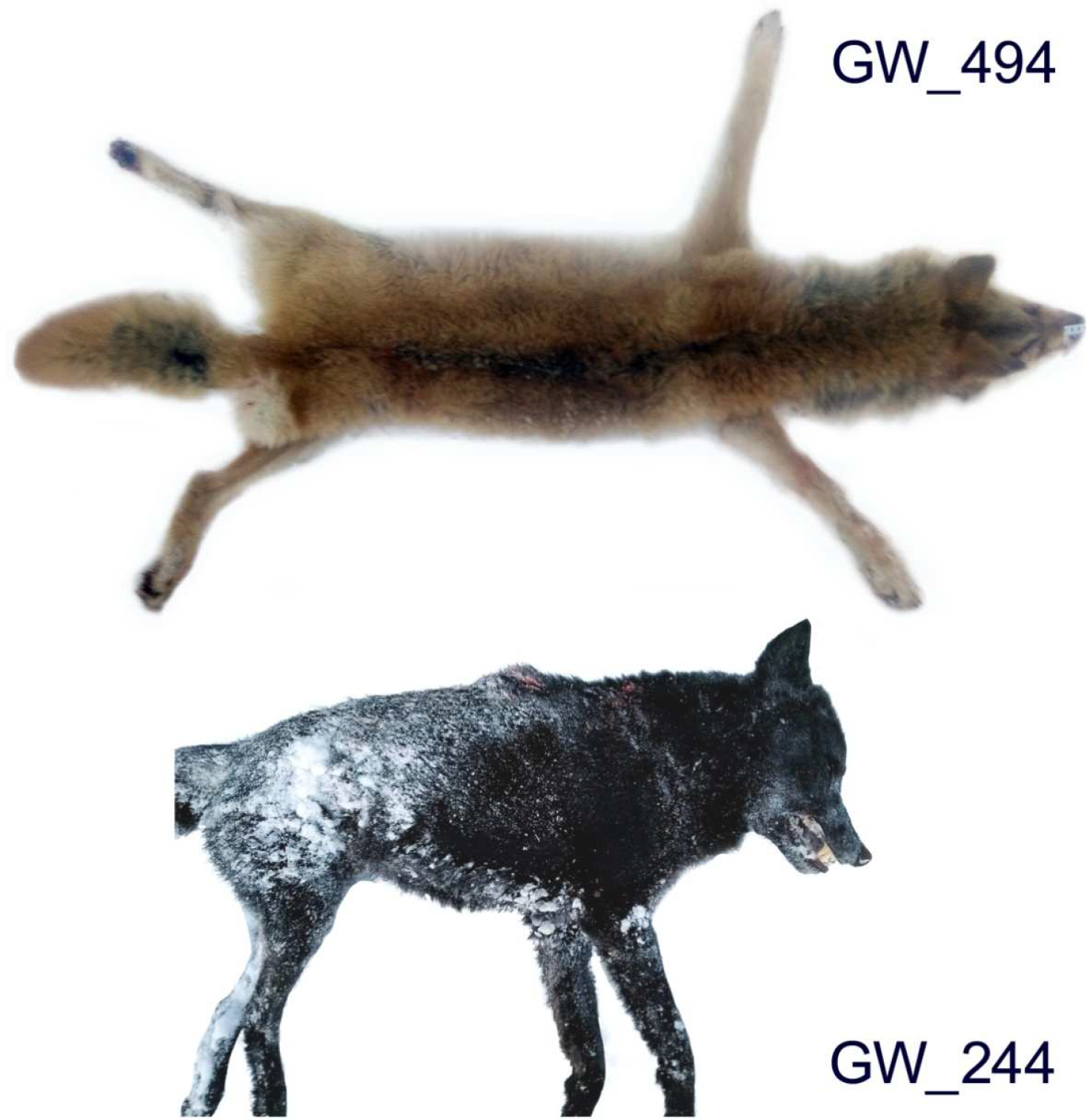
The skin of the suspected wolf-dog hybrid GW_494 but genetically recognized as pure wolf, and general appearance of the wolf-dog hybrid GW_244 (photos provided by local hunting authorities)

### CR1 mtDNA

Sequences of the mtDNA control region were obtained for 36 individuals from the wild wolf population (35 wolves and 1 feral dog). Three haplotypes were revealed, the number of polymorphic sites was 8, haplotype diversity Hd = 0.541±0.034, and nucleotide diversity P = 0.00854±0.00060. In the sample set, 18 wolves had the haplotype with GenBank accession number OP503597 (here and further #1) and 16 had the haplotype OP503598 (#2), both were previously known for RK and common in that region (Tirronen et al. 2023). Two samples had a third haplotype often found in domestic dogs (*Canis familaris*) and rare in wolves (Ersmark et al. 2016). It was sample GW_246 marked as a “hybrid” and the feral dog GW_50. Information about the haplotypes of suspected and identified hybrids is presented in Table 1.

**Table 1.** Identifications of probabilities for phenotypic and putative hybrids found in the wolf population of Russian Karelia.

| Sample | Year | Location | mtDNA<br>haplotype | Phenotypic trait | STRUCTURE | NEWHYBRIDS |  |  |  |  |  |
| --- | --- | --- | --- | --- | --- | --- | --- | --- | --- | --- | --- |
|  |  |  |  |  | q_wolf | Wolf | Dog | F1 | F2 | BxW | BxD |
| GW_91 | 2017 | 61°19'00" N<br>29°28'22" E | 2 | Nothing unusual, “wolf<br>type” | 0.9008 | 0.32683 | 0.00000 | 0.00006 | 0.06681 | 0.60628 | 0.00002 |
| GW_119 | 2018 | 62°55'44" N<br>34°28'55" E | 1 | White claws, white<br>pads | 0.7960 | 0.00986 | 0.00000 | 0.00571 | 0.08911 | 0.89505 | 0.00028 |
| GW_196 | 2020 | 62°30'54" N<br>34°14'34" E | 2 | Nothing unusual, “wolf<br>type” | 0.9350 | 0.31410 | 0.00000 | 0.00149 | 0.03718 | 0.64708 | 0.00015 |
| GW_244 | 2021 | 61°19'47" N<br>29°26'10" E | 2 | Completely black color | 0.9234 | 0.81968 | 0.00000 | 0.00040 | 0.01664 | 0.16312 | 0.00016 |
| GW_246 | 2021 | 63°47'N<br>31°39'E | dog | Marked as a “WDH”<br>(without details) | 0.4856 | 0.00055 | 0.00001 | 0.01290 | 0.25360 | 0.71972 | 0.01323 |
| GW_494 | 2022 | 64°16'30"N<br>34°05'10"E | 1 | No black tip on tail and<br>black stripes on front<br>legs | 0.9940 | 0.99273 | 0.00000 | 0.00000 | 0.00007 | 0.00719 | 0.00000 |

### Genotyping errors

All the samples were genotyped using 20 microsatellite markers, but only for 14 loci it was possible to obtain reliable allele length. Six loci were excluded from further analysis due to the appearance of false alleles. The MICRO-CHECKER program analyzed 36 genetic profiles for genotyping errors and identified 5 loci (FH2096, CXX279, C09.173, CPH5, FH2054) with heterozygosity deficiency suggesting the presence of null alleles in these loci. These loci were excluded from the wolf-dog hybridization analysis if they also showed significant deviation from HWE. Four loci (CPH2, CXX279, CPH5, FH2054) were significantly out of HWE (P ≤0.005) in the pooled data set of 36 samples. As a result, the hybridization of wolves and dogs in the Karelian wolf population was assessed using a set of 11 loci: AHT137, CPH2, CPH8, CPH4, CPH12, FH2010, FH2096, AHTk211, vWF, FH2001, C09.173 (Ostrander et al. 1993; Shibuya et al. 1994; Holmes et al. 1995; Fredholm and Wintero 1995; Francisco et al. 1996; Lingaas et al. 1997). All loci were used to assess the genetic diversity of the wolf population and its demographic history.

### Genetic diversity of the wolf population

All microsatellite loci were polymorphic and the diversity was relatively high: Ho = 0.655 (SE = 0.022), He = 0.746 (SE = 0.029), A = 8.1 (SE = 0.835) (Table 2). The overall inbreeding coefficient in the studied population was positive and significantly differed from zero (F = 0.131, P = 0.0000). Also, the inbreeding coefficient was significantly positive in five loci (CPH4, CXX279, C09.173, CPH5, FH2054). The STRUCTURE analysis of 35 samples of wolves and 12 of dogs by 11 autosomal STRs reveal a clear genetic differentiation into two separate groups. An optimal clustering of samples was obtained with K = 2, with a few intermediate individuals of mixed ancestry (Fig. 3). The average q_wolf_ = 0.990, q_dog_= 0.988.

**Table 2.** Genetic diversity. Expected (He) and observed (Ho) heterozygosities, number of alleles (A), and inbreeding coefficients (Fis) of 14 microsatellite loci in the RK wolf population (n = 35)

| Locus | He | Ho | A | Fis |
| --- | --- | --- | --- | --- |
| AHT137 | 0.872 | 0.828 | 10 | 0.068 |
| CPH2 | 0.744 | 0.633 | 8 | 0.165 |
| CPH8 | 0.763 | 0.655 | 8 | 0.158 |
| CPH4 | 0.794 | 0.667 | 6 | 0.177* |
| CPH12 | 0.632 | 0.700 | 7 | -0.091 |
| FH2010 | 0.701 | 0.633 | 5 | 0.113 |
| FH2096 | 0.760 | 0.690 | 5 | 0.110 |
| AHTk211 | 0.441 | 0.464 | 4 | -0.035 |
| vWF | 0.769 | 0.700 | 7 | 0.107 |
| FH2001 | 0.711 | 0.733 | 5 | -0.015 |
| CXX279 | 0.865 | 0.667 | 14 | 0.245* |
| C09.173 | 0.802 | 0.633 | 11 | 0.226* |
| CPH5 | 0.814 | 0.567 | 13 | 0.319* |
| FH2054 | 0.776 | 0.600 | 10 | 0.246* |
| Mean | 0.746 | 0.655 | 8.1 | 0.131 |
| SE | 0.029 | 0.022 | 0.835 |  |

\* $p < 0.05$

**Fig. 3.**
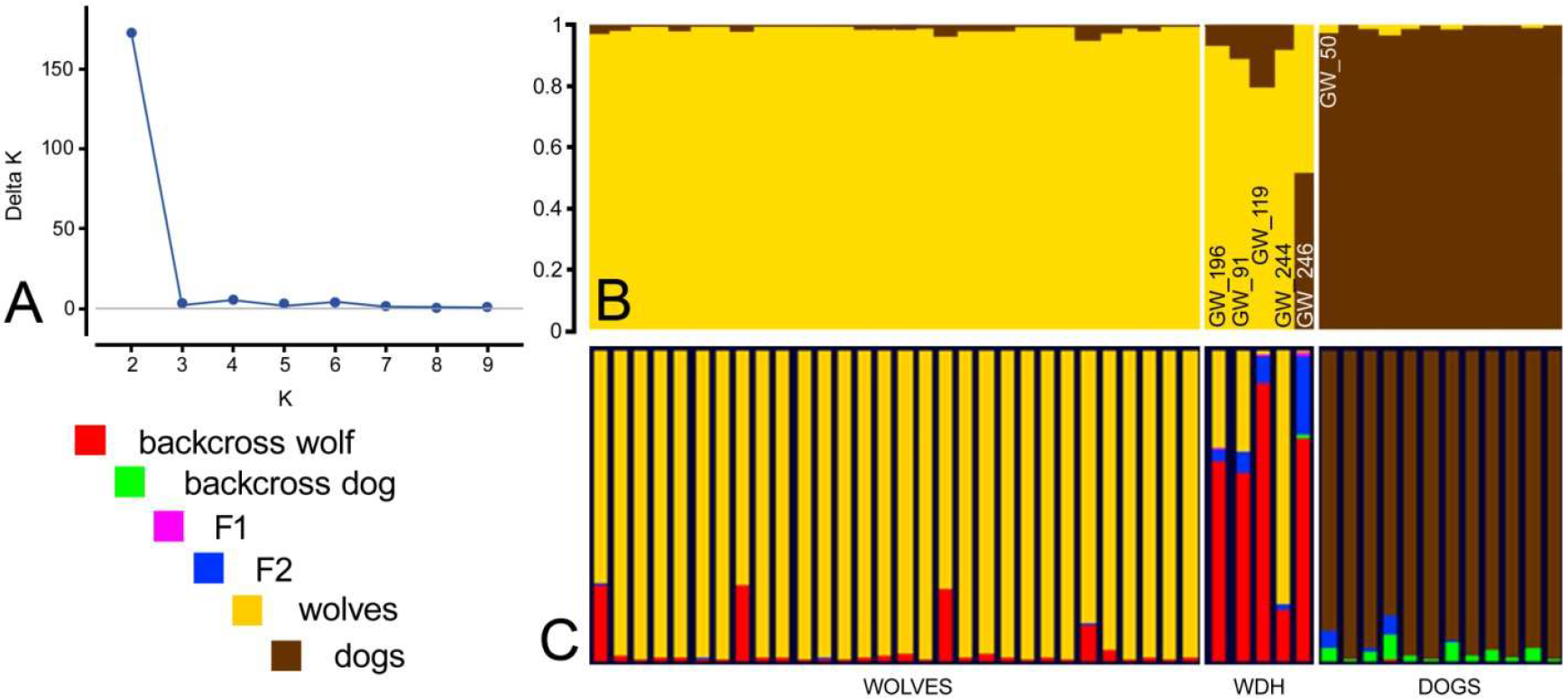
Genetic differentiation of wolves, dog, and WDHs: (A) K values (Evanno et al. 2005); bar plotting outputs of the analysis from STRUCTURE (B) and NEWHYBRIDS (C)

### Population bottleneck

The Wilcoxon’s test for heterozygosity excess was performed for wolf samples excluding hybrids in two repeat using 11 and 14 microsatellite markers, revealing no bottlenecks in either case (P = 0.681 and P = 0.914). Also, the allelic frequency distribution for the Karelian population was L-shaped in both cases as expected for populations that have not gone through recent bottlenecks. We obtained similar results using 11 as well as 14 microsatellite loci, which agrees with the results of J. Carlsson (2008) and N. Sastre et al. (2011), who found no differences in using genetic data with or without null alleles.

### Genetic identification of WDHs

Clustering using Bayesian approaches in STRUCTURE software was performed on the entire data set for 11 loci with the number of inferred clusters increasing from K=1 to K=10. ΔK analysis showed a high peak when K=2 and no peaks for the rest of K (Fig. 3).

The genetic profiles were clearly divided according to their ancestry into the wolf and dog groups with the average q_wolf_ = 0.990 (N=30) and q_dog_= 0.988 (N=12). The Q-value for pure wolves ranged from 0.960 to 0.996. Five individuals were of mixed ancestry with q_wolf_ ranging between 0.486 (GW_246) and 0.935 (GW_196), and they were identified as WDH. They accounted for 14.3% of the wolf sample. A threshold of q_wolf_ ≥ 0.935 was established to distinguish hybrids from pure wolfs. Only for one sample, GW_246, F1 generation can be presumed with 49% probability (Table 1).

The results of the NEWHYBRIDS analysis (Fig. 3, Table 1) confirmed the results of the STRUCTURE. Individuals (except for GW_244) were assigned to classes of origin (wolf, dog, or hybrids). GW_244 was classified as a wolf, but with a 16% probability of backcross to wolf. The hybrid sample GW_246 was determined to be a backcross wolf (72%) or F2 (25%). The rest of the samples (GW_91, GW_119, GW_196) were assigned to first generation backcrosses to wolf (Table 1).

In the FCA analysis, wolves, dogs and five hybrids were clearly distinguished from each other based on the distributions of allele frequencies at 11 microsatellite loci. The sample GW_243 departed somewhat from the pure wolf cloud (Fig. 4). In a previous analysis it was not detected as a hybrid. The black wolf GW_244 identified as a hybrid was taken together with GW_243 from the same pack. It was established that these wolves were not related. The maximum likelihood estimate of relatedness was 0.0072.

**Fig. 4.**
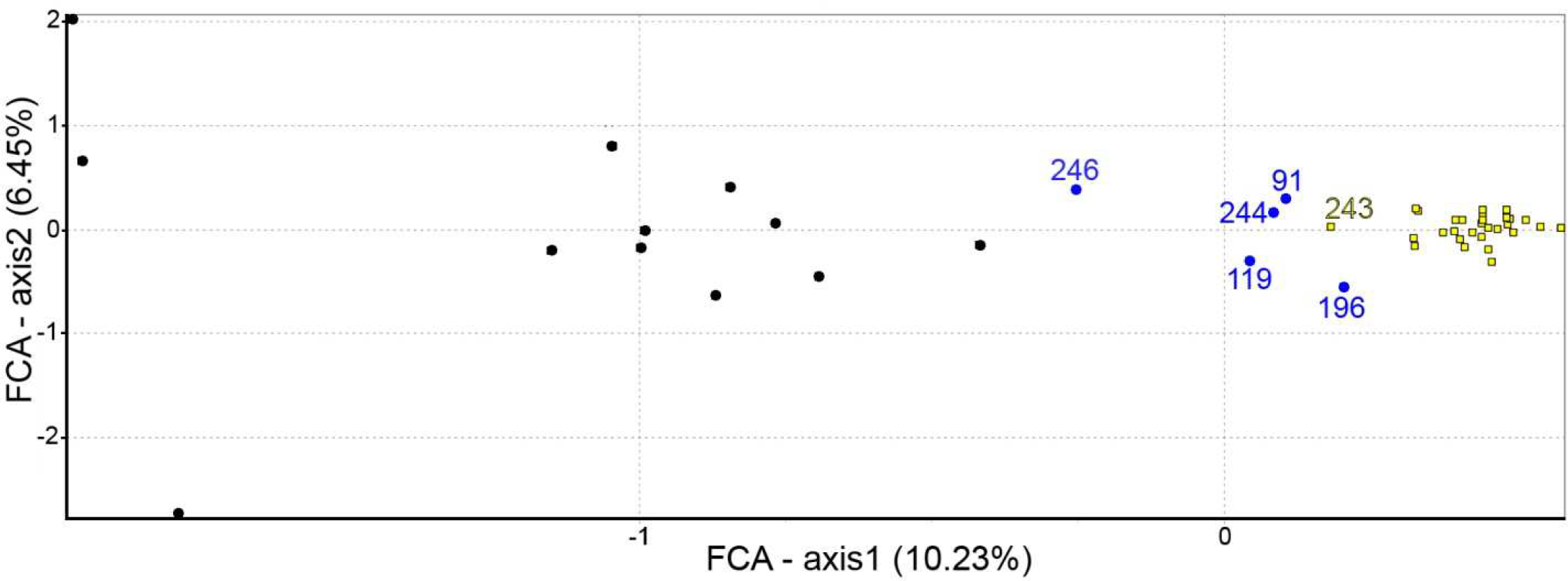
FCA by 11 STRs illustrating the genetic distribution of dogs (black dots), wolves (yellow squares), and hybrids (blue dots)

Of the four individuals who had phenotypic deviations from the wolf normal appearance or were marked as a “hybrid”, in two cases (GW_119, GW_244) the hybrid status was confirmed only by STR, and in one case (GW_246) by STR and mtDNA analyses. The GW_494 individual with a dog phenotype was surprisingly recognized as a pure wolf by both marker classes (Table 1, Fig 3). For this sample (GW_494), we repeated all stages of analysis since DNA extraction three times. Genetic analysis of microsatellite markers revealed two additional WDHs: sample GW_91 and GW_196, which had no phenotypic deviations from the “wolf”.

## DISCUSSION

Discussing the results, we will often refer to the Finnish wolf population and here is why:

1. Biogeographically, the “Karelian” and “Finnish” populations represent a single population without geographical barriers, which continues further to the east and south, and its division into two populations is rather nominal (Fig. 1) (Boitani 2003; Kojola et al. 2009);
2. The landscape and climatic conditions and approaches to wolf management in Finland and Russian Karelia are completely different, which affects the dynamics of these sub-populations (Suutarinen and Kojola 2017; Tirronen et al. 2023);
3. Attempts have previously been made to assess the level of differentiation between these populations using current sets of autosomal STRs. Thus, J. Aspi et al. (2009) using 10, and E. Jansson et al. (2014) using 17 microsatellite markers found that the Finnish and Karelian populations were differentiated, whereas M. Åkesson et al. (2022) using a set of 26 autosomal STRs did not reveal significant differentiation between the populations.

There are no phenotype-based reports in the literature of hybrid individuals among RK wolves (Danilov 2005; Danilov et al. 1979). No WDHs were revealed in previous microsatellite-based wolf population genetic studies either (Aspi et al. 2009; Jansson et al. 2012, 2014). The sample set of RK wolves used for these genetic studies amounted to 37 specimens covering the periods 1995-2000 and 2009-2010. The whole-genome study of Fennoscandian wolf populations did not detect WDHs among 15 samples of RK wolves in the 2010s either (Smeds et al. 2020). The analysis of CR1 mtDNA in the sample of 90 RK wolves covering the period 2012-2022 detected an animal with dog ancestry for the first time (Tirronen et al. 2023). However, the applicability of this method for reliable WDH detection is limited by the maternal inheritance of mtDNA (Vilà and Wayne 2001).

The WDH could appear due to a combination of a disruption of the normal population structure of wolves and decrease in the wolf population density with the presence of stray or feral dogs, and even with deliberate hybridization. Regular and intensive wolf hunting could lead to a break-down of the social structure, rotation of breeding individuals, changes in the breeding behavior, lowering of reproduction success and survival of yearlings, inbreeding, and hybridization (Ryabov 1973; Jędrzejewska et al. 1996; Brainerd et al. 2008; Liberg et al. 2005).

During the period from 2017 to 2022, four wolves (GW_119, GW_244, GW_246, GW_494) with phenotypic deviations suggesting WDH were removed from the Karelian wolf population, among others. In three cases (GW_119, GW_244, GW_246) this inference was confirmed genetically. Two more individuals (GW_91 and GW_196) were also identified as wolf-dog hybrids during the genetic analysis of a random sample, but they did not have phenotypic features distinguishing them from wolves. This brings us to the important and previously repeatedly stated conclusion that phenotypic traits are not always sufficient to identify a hybrid individual and validation by genetic methods is required. The dog mtDNA haplotype in GW_246 indicates this WDH appeared as a result of a crossing between female dog and male wolf. Such an event of interspecific crossing is interesting and rare. There is a typical sexual disproportion in the hybridization of wolves and dogs in the wild, where a lone female wolf chooses a male dog as a pair (Hendrickson et al. 2012). The GW_246 sample comes from the northern taiga subzone of Karelia, where the wolf population density is relatively low. It seems that the combination of a sparse population and intensive human persecution were sufficient for a rare hybrid type (more likely F2) to appear (Table 1, Fig. 3). Another hybrid of unusual appearance – GW_244, was strikingly different from the average Karelian wolf in having a totally black coat. The common color of Karelian wolves is a variety of shades of light and dark gray, often in combination with reddish color, generating a color variation from almost straw to dark gray, but black wolves have never been found there (Danilov 2005). It is known that black wolves are common in some populations of North America, but the emergence of such individuals in European populations is associated with possible hybridization with dogs (Gipson et al. 2002; Musiani et al., 2007; Caniglia et al. 2013). Interestingly, the STRUCTURE simulation for this individual yielded q_wolf_ = 0.923, which is extremely close to the threshold value established in our study to distinguish pure wolves from hybrids (q_wolf_ ≥ 0.935), and NEWHYBRIDS with a 16% probability defines it as a backcross wolf. Obviously, interspecific hybridization in this case took place several generations ago and a wider set of STRs or SNPs is needed to confidently determine that instant. In general, the use of functional mutations in combination with standard non-coding molecular markers can additionally help detect hybridization and introgression of genes in wild canine populations (Caniglia et al. 2013). An interesting result was also obtained in the case of GW_494: for the set of markers we used both STRUCTURE and NEWHYBRIDS uniquely identify this individual as a pure wolf, despite the clear phenotypic signs of a hybrid. Theoretically, the loss of distinctive phenotypic traits of wolves may be a result of a more distant dog introgression, and the resolution of 11 microsatellite markers is insufficient for its identification. For example, J. Kusak et al. (2018) note that 12 microsatellite STRs were not informative enough for identification of the generations of admixture, they only determined whether the identified hybrids were a backcross with wolves, a backcross with dogs, or a wolf with a past dog gene introgression. The difficulties in determining the classes of hybrids using a similar set of microsatellite markers were described in detail by Korablev et al. (2021). In general, the application of STRUCTURE and NEWHYBRIDS in our study produced comparable results, clearly distinguishing wolves from dogs, and mixed individuals. However, the results do not provide grounds for a definite judgement regarding the status of the mixed individuals. To uniquely identify wolves, dogs, and their hybrids using autosomal STRs, Lorenzini et al. (2022) proposed a multiplex panel of 22 microsatellite markers. J. Harmoinen et al. (2021) suggest using a 96 SNP panel, arguing that these new markers do not require calibration among laboratories, are capable of detecting wolf-dog hybrids up to third-generation backcrosses to wolves, and demonstrate a high genotyping success rate for all sample types. The difficulties we encountered in our study have also convinced us of the need to unify the methods of genetic identification of wolves and hybrids implemented in species monitoring programs, especially when it comes to transboundary populations.

In the studied population, we observe fairly high rates of genetic polymorphism, comparable to those noted earlier in Karelia: Ho = 0.656, He = 0.709, and A = 5.7 (Aspi et al. 2009), Finland: Ho = 0.712, He = 0.697, and A = 5.9 (Jansson et al. 2014), central part of European Russia: Ho = 0.79, He = 0.80, and A = 10 (Korablev et al. 2021), and on average for some regions of Siberia: Ho = 0.695, He = 0.649 (Talala et al. 2020). It is clear that there can be no direct comparison of these indexes because of the different sample sizes and sets of microsatellite markers in each single study. However, the relatively high level of heterozygosity in wolves from the RK does not mean the population is in good state. It may be owing to the specific transit position of the territory, the absence of natural barriers, constant contacts with neighboring populations, diversity of the initial population, but at the same time there may be a possible loss of allelic and mitochondrial diversity (Jędrzejewski et al. 2005; Salado et al. 2022). In Finland, e.g., about 20% of microsatellite alleles were reportedly (Jansson et al. 2014) not found in the modern population in comparison with historical samples. The heterozygosity indexes in the studied population demonstrate that the observed heterozygosity Ho = 0.655 is significantly (p < 0.05) less than the expected He = 0.746, likely indicating a prevalence of inbreeding. This is corroborated by the coefficient Fis = 0.131, which is higher compared to the indexes previously noted for Karelia in 1995-2000 – Fis = 0.094 (Aspi et al. 2009) and in 1997-2000 – Fis = 0.063 (Åkesson et al. 2022). In Finland, the inbreeding coefficient was also slightly lower in 2007-2009 Fis = 0.108 (Jansson et al. 2012). The Fis value is quite similar to the corresponding index for some isolated European wolf populations (Lucchini et al. 2004; Sastre et al. 2011).

Nowadays, more than 60 mitochondrial DNA haplotypes (depending on the length of fragments) have been detected for the Eurasian wolf (Ersmark et al. 2016; Nechaeva et al. 2022). The reduction in diversity from 8 historical to 3 modern mtDNA haplotypes was found in Finland (Jansson 2013). Currently, only two mtDNA haplotypes have been noted in Karelia, and this level of diversity corresponds to small and isolated populations (Randi et al. 2001; Tirronen et al. 2023). We supposed that the low modern diversity of mtDNA haplotypes in the RK wolf population may be due to the species’ history – periods of intensive hunting in the past and significant population declines. However, neither our study nor previous studies (Aspi et al. 2009) have detected a bottleneck effect.

Some of the key parameters for the wolf population dynamics in Karelia for the periods I (1992-2000), II (2001-2011), III (2012-2022) are the following: average annual numbers 528, 380 and 330 animals; average annual harvest rates 146, 77 and 156 individuals; total numbers of wolves killed 1.312, 847 and 1.718, respectively. The two-fold increase in wolf harvest in the third vs the second period affected not only the abundance dynamics (for more information, see Tirronen et al. 2023) but also the genetic structure and diversity of the population. Apparently, a lack of available partners may lead to both close-relative breeding and interspecific hybridization. Due to intensive hunting, the Karelian population may experience a shortage of breeding individuals, and a sort of “vacuum” is formed over vast areas, which can be filled by dispersal from the west by Finnish, and from the east by Arkhangelsk wolves. Theoretically, this process can make these two populations genetically closer, although it is difficult to predict how the management of the wolf population in these regions may change in the future, which will largely define the structure of these populations.

One of the key points of this study is the sudden appearance, within a relatively short period of time, of several WDHs at once in a population that was previously considered “pure”. Importantly, the relationships between wolves and dogs in the area occupied by the Finnish-Karelian wolf population used to be exclusively within the predator-prey coordinates (Tikkunen and Kojola 2020; Tirronen 2008). An important question arises – does the detected ratio of pure wolves and hybrids accurately represent the real situation in the studied population? In our sample set of 35 individuals, four were not random (phenotypically suspected hybrids), so we can ignore them. However, the genetic analysis of the remaining random sample (n = 31) revealed two more WDHs, i.e. 6.5% of the total. We shall not directly extrapolate this figure to the entire sample of wolves harvested during this period (n = 1718), considering the statistical power of the sample we examined to be insufficient (n = 31). This proportion is similar to that in wolf populations in human-dominated landscapes and where man actively interferes in the natural history of wild species (Pacheco et al. 2017; Hindrikson et al. 2012). Our study can be regarded as empirical evidence supporting the hypothesis that wolves engage interspecific hybridization as a behavioral adaptive mechanism securing the species survival under extremely unfavorable conditions (Bohling and Waits 2015; Adavoudi and Pilot 2021).

In undisturbed populations, the core of the population is formed by a family pack, in which usually only the alpha pair participates in breeding. In the case of a hunted population, individuals of different ages, social status, and even close kinship can breed (Ballard et al. 1987; Mech 1999; Fuller et al. 2003; Yudin 2013). However, even this induced response also has its biologically determined limits, a striking example provided by the observation of wolves on Isle Royale, where a small and isolated population at a certain point in its existence becomes unable to overcome a complex of intraspecific restrictions and consequences caused by an inbreeding depression (Hedrick et al., 2019). A rise in inbreeding has serious demographic consequences for the population, affects resistance to parasites and infectious diseases (Liberg et al. 2005; Niskanen et al. 2014). In our case, it is quite possible that the low diversity of mtDNA haplotypes, an increase in the proportion of inbreeding, and the appearance of interspecific hybrids with dog are consequences of the intensified wolf hunting pressure. As a result, wolves mate both with closely related individuals and in some cases with dogs. One way or another, this study is not just a first attempt to identify hybrids in the Karelian wolf population, but also a valid inquiry into species’ response to direct and intensive human persecution. In the future, large-scale research, including genomic cross-border studies, will need to answer practical questions about the status of the population and will be extremely important from the fundamental standpoint of studying the processes of microevolution and adaptive capabilities of the species.

## ACKNOWLEDGMENTS

We appreciate the support of the Ministry of Natural Resources and Environment of the Republic of Karelia in collecting wolf samples. We are grateful to the animal shelter and personally to Vladimir Rybalko for providing dog samples. We thank Ilpo Kojola for sharing information on black wolves in a Finnish pack.

## LITERATURE

Åkesson M, Flagstad Ø, Aspi J, et al. (2022) Genetic signature of immigrants and their effect on genetic diversity in the recently established Scandinavian wolf population. Conserv Genet 23:359–373. 10.1007/s10592-021-01423-5

Anderson EC, Thompson EA (2002) A model-based method for identifying species hybrids using multilocus genetic data. Genetics 160:1217–1229. 10.1093/genetics/160.3.1217.

Anderson TM, Vonholdt BM, Candille SI et al. (2009). Molecular and evolutionary history of melanism in North American gray wolves. Science 323:1339–1343. 10.1126/science.1165448

Apollonio M, Mattioli L, Scandura M (2004) Occurrence of black wolves in the Northern Apennines, Italy. Acta Theriol 49:281–285. 10.1007/BF03192528

Aspi J, Roininen M, Kiiskila J, Ruokonen M, Kojola I, Bljudnik L, Danilov P, Heikkinen S, Pulliainen E (2009) Genetic structure of the northwestern Russian wolf populations and gene flow between Russia and Finland. Conserv Genet 10:815–826. 10.1007/s10592-008-9642-x

Adavoudi R, Pilot M (2021) Consequences of Hybridization in Mammals: A Systematic Review. Genes 13(50). 10.3390/genes13010050.

Belkhir K, Borsa P, Chikhi L, Raufaste N, Bonhomme F (2004) GENETIX 4.05, logiciel sous Windows TM pour la génétique des populations. Laboratoire génome, populations, interactions. CNRS UMR 5000. Université de Montpellier II, France. https://kimura.univmontp2.fr/genetix/ Bibikov DI (ed) (1985) Wolf: Origin, Systematics, Morphology, and Ecology. Nauka, Moscow. (in Russian)

Bohling J, Waits L (2015) Factors influencing red wolf–coyote hybridization in eastern North Carolina, USA. Biol Conserv 184. 10.1016/j.biocon.2015.01.013.

Boitani L (2003) Wolf conservation and recovery. In: Wolves. Behavior, Ecology and Conservation (eds Mech LD, Boitani L), University of Chicago Press. 317–340.

Ballard WB, Whitman JS, Gardner CL (1987) Ecology of an exploited wolf population in south-central Alaska. Wildl Monogr 98.

Brainerd SM, Andren H, Bangs EE et al. (2008) The effects of breeder loss on wolves. J Wildl Manage. 72:89–98. 10.2193/2006-305

Carlsson J (2008) Effects of microsatellite null alleles on assignment testing. J Hered 99:616–623. 10.1093/jhered/esn048

Cornuet JM, Luikart G (1996) Description and power analysis of two tests for detecting recent population bottlenecks from allele frequency data. Genetics 144:2001–2014. 10.1093/genetics/144.4.2001

Danilov PI, Rusakov OS, Tumanov IL (1979) Predatory Animals of the North-West of the USSR. Leningrad: Nauka. (in Russian)

Danilov PI (2005) Game Animals of Karelia: Ecology, Resources, Management, and Conservation. Moscow: Nauka. (in Russian)

Earl DA, Vonholdt B (2012) Structure Harvester: a website and program for visualizing STRUCTURE output and implementing the Evanno method. Conserv Genet Res. 4. 10.1007/s12686-011-9548-7

Ersmark E, Klütsch C, Chen Y, et al. (2016) From the past to the present: Wolf phylogeography and demographic history based on the mitochondrial control region. Front Ecol and Envir 4:134. 10.3389/fevo.2016.00134

Excoffier L, Lischer HEL (2010) Arlequin suite ver 3.5: a new series of programs to perform population genetics analyses under Linux and Windows. Mol Ecol Res 10:564–567. 10.1111/j.1755-0998.2010.02847.x

Dziech A (2021) Identification of Wolf-Dog Hybrids in Europe - An Overview of Genetic Studies. Front Ecol Evol. 9. 10.3389/fevo.2021.760160

Evanno G, Regnaut S, Goudet J (2005) Detecting the number of clusters of individuals using the software Structure: a simulation study. Mol Ecol 14:2611–2620. 10.1111/j.1365-294X.2005.02553.x

Francisco LV, Langston AA, Mellersh CS, Neal CL, Ostrander EA (1996) A class of highly polymorphic tetranucleotide repeats for canine genetic mapping. Mamm Gen 7:359–362. 10.1007/s003359900104

Fredholm M, Wintero AK (1995) Variation of short tandem repeats within and between species belonging to the Canidae family. Mamm Genome 6:11–18. 10.1007/BF00350887

Fuller TK, Mech LD, Cochrane JF (2003) Wolf population dynamics. Wolves: Behavior, Ecology and Conservation (eds. Mech LD, Boitani L)Chicago, Illinois

Caniglia R, Fabbri E, Greco C. et al. (2013) Black coats in an admixed wolf × dog pack is melanism an indicator of hybridization in wolves? Eur J Wildl Res 59:543–555. 10.1007/s10344-013-0703-1

Gipson P, Bangs E, Bailey T, Boyd D, Cluff H, Smith D, Jiminez M (2002) Color Patterns among Wolves in Western North America. Wildl Soc Bull. 30:821–830. 10.2307/3784236

Harmoinen J, von Thaden A, Aspi J, et al. (2021) Reliable wolf-dog hybrid detection in Europe using a reduced SNP panel developed for non-invasively collected samples. BMC Genomics 22, 473 10.1186/s12864-021-07761-5

Hedrick P (2009) Wolf of a different colour. Heredity 103:435–436. 10.1038/hdy.2009.77

Hedrick PW, Robinson JA, Peterson RO, Vucetich JA (2019) Genetics and extinction and the example of Isle Royale wolves. Anim Conserv 22:302–309. 10.1111/acv.12479

Hindrikson M, Männil P, Ozolins J, Krzywinski A, Saarma U (2012) Bucking the Trend in Wolf-Dog Hybridization: First Evidence from Europe of Hybridization between Female Dogs and Male Wolves. PLoS ONE 7(10): e46465. doi:10.1371/journal.pone.0046465

Hindrikson M, Remm J, Pilot M, Godinho R, et al. (2016). Wolf population genetics in Europe: A systematic review meta-analysis and suggestions for conservation and management. Biol Rev 92:1601–1629. 10.1111/brv.12298

Holmes NG, Dickens HF, Parker HL, Binns MM, Samson J (1995) Eighteen canine microsatellites. Animal Genet 262:132–133. 10.1111/j.1365-2052.1995.tb02659.x.

Jakobsson, M. & Rosenberg, N. A. (2007) CLUMPP: a cluster matching and permutation program for dealing with label switching and multimodality in analysis of population structure. Bioinformatics 23: 1801–1806. 10.1093/bioinformatics/btm233

Jansson E, Ruokonen M, Kojola I, and Aspi J (2012) Rise and fall of a wolf population: genetic diversity and structure during recovery, rapid expansion and drastic decline. Mol Ecol 21:5178–5193. 10.1111/mec.12010

Jansson E. (2013) Past and present genetic diversity and structure of the Finnish wolf population. PhD Thesis.

Jansson E, Harmoinen J, Ruokonen M, Aspi J (2014) Living on the edge: Reconstructing the genetic history of the Finnish wolf population. BMC Evol Biol 14. 64. 10.1186/1471-2148-14-64

Jędrzejewska B, Jędrzejewska W, Bunevich AN, Miłkowski L, and Okarma H (1996) Population dynamics of wolves Canis lupus in Białowieza a primeval forest (Poland and Belarus) in relation to hunting by humans, 1847–1993. Mamm Rev 26:103–126. 10.1111/j.1365-2907.1996.tb00149.x

Kalinowski ST, Wagner AP, Taper ML (2006) ml-relate: a computer program for maximum likelihood estimation of relatedness and relationship. Mol Ecol Notes 6:576–579. 10.1111/j.1471-8286.2006.01256.x

Kojola I, Kaartinen S, Hakala A, Heikkinen S, Voipio H-M (2009) Dispersal behavior and the connectivity between wolf populations in Northern Europe. J Wildl Manag 73:309–313. 10.2193/2007-539

Korablev MP, Korablev NP, Korablev PN (2021) Genetic diversity and population structure of the grey wolf (Canis lupus Linnaeus, 1758) and evidence of wolf × dog hybridisation in the centre of European Russia. Mamm Biol 101:1–104 10.1007/s42991-020-00074-2

Kusak J, Fabbri E, Galov A, Gomerčić T, Arbanasić H, Caniglia R, Galaverni M, Reljic S, Huber D, Randi E (2018) Wolf-dog hybridization in Croatia. Vet arhiv. 88:375–395. 10.24099/vet.arhiv.170314

Liberg O, Andrén H, Pedersen H, et al, (2005) Severe inbreeding depression in a wild wolf (Canis lupus) population. Biol lett. 1:17–20. 10.1098/rsbl.2004.0266.

Librado P, Rozas J (2009) DnaSP v5: a software for comprehensive analysis of DNA polymorphism data. Bioinformatics. 25(11): 1451–1452. 10.1093/bioinformatics/btp187.

Lingaas F, Sorensen A, Juneja RK et al. (1997) Towards construction of a canine linkage map: Establishment of 16 linkage groups. Mammal Gen 8:218–221. 10.1007/s003359900393

Lorenzini R, Attili L, Tancredi C, Fanelli R, Garofalo L (2022) A Validated Molecular Protocol to Differentiate Pure Wolves, Dogs and Wolf x Dog Hybrids through a Panel of Multiplexed Canine STR Markers. Divers 14, 511. 10.3390/d14070511

Lucchini V, Galov A, Randi E (2004) Evidence of genetic distinction and long-term population decline in wolves (Canis lupus) in the Italian Apennines. Mol ecol 13:523–36. 10.1046/j.1365-294X.2004.02077.x.

Luikart G, Cornuet JM (1997) Empirical evaluation of a test for identifying recently bottlenecked populations from allele frequency data. Conserv Biol 12:228–237. 10.1111/j.1523-1739.1998.96388.x

Mariat D, Kessler JL, Vaiman D, Panthier JJ (1996) Polymorphism characterization of five canine microsatellites. Animal Genet 27:434–435. 10.1111/j.1365-2052.1996.tb00514.x.

Mech LD (1999) Alpha status, dominance, and division of labor in wolf packs. Canad J Zool 77:1196–1203. 10.1139/z99-099

Molchan V, Homel K, Valnisty A, Nikiforov M, Theodorova E (2023) Genetic diversity of mtDNA in the grey wolf population of Belarus threatened by wolf-dog admixture. Theriol Ukr 25:87–99. 10.53452/TU2508

Musiani M, Leonard JA, Cluff HD et al. (2007) Differentiation of tundra/taiga and boreal coniferous forest wolves: genetics, coat colour and association with migratory caribou. Mol Ecol 16:4149–4170. 10.1111/j.1365-294X.2007.03458.x

Nechaeva AV, Belokon’ MM, Belokon’ YuS, et al. (2022) Genetic diversity of Canis lupus L. in Eastern Europe based on mitochondrial data (Proc Conf “Genetic Processes in Populations,” Moscow, October 11–14, 2022), Moscow: Vash Format, p. 44. (in Russian)

Niskanen A, Kennedy L, Ruokonen M, Kojola I, Lohi H, Isomursu M, Jansson E, Pyhäjärvi T, Aspi J (2014). Balancing selection and heterozygote advantage in major histocompatibility complex loci of the bottlenecked Finnish wolf population. Mol Ecol 23:875–889. 10.1111/mec.12647.

Ostrander EA, Mapa FA, Yee M, Rine J (1995) One hundred and one simple sequence repeats-based markers for the canine genome. Mamm Gen 6:192–195. 10.1007/BF00293011.

Pacheco C, López-Bao JV, García EJ, Lema FJ, Llaneza L, Palacios V, Godinho R (2017) Spatial assessment of wolf–dog hybridization in a single breeding period. Sci Rep 7:42475. 10.1038/srep42475

Peakall R, Smouse PE (2012) GenAlEx 6.5: genetic analysis in Excel. Population genetic software for teaching and research – an update. Bioinform 28:2537–2539. 10.1093/bioinformatics/bts460

Pilot M, Jedrzejewski W, Branicki W, Sidorovich V, Jedrzejewska B, Stachura K, Funk S (2007) Ecological factors influence population genetic structure of European grey wolves. Mol ecol 15. 10.1111/j.1365-294X.2006.03110.x

Pilot M, Greco C, Vonholdt B, Randi E, Jedrzejewski W et al. (2018) Widespread long term admixture between grey wolves and domestic dogs across Eurasia and its implications for the conservation status of hybrids. Evol Appl 11. 10.1111/eva.12595

Pilot M, Moura AE, Okhlopkov IM, Mamaev NV et al. (2019) Global Phylogeographic and Admixture Patterns in Grey Wolves and Genetic Legacy of An Ancient Siberian Lineage. Sci Rep 9:17328 10.1038/s41598-019-53492-9

Piry S, Luikart G, Cornuet J-M (1999) BOTTLENECK: a computer program for detecting recent reductions in the effective population size using allele frequency data. J Hered 90(4). 10.1093/jhered/90.4.502

Pritchard JK, Stephens M, Donnelly P (2000) Inference of population structure using multilocus genotype data. Genetics 155:945–959. 10.1093/genetics/155.2.945

Ramasamy RK, Ramasamy S, Bindroo BB, et al. (2014) STRUCTURE PLOT: a program for drawing elegant STRUCTURE bar plots in user friendly interface. SpringerPlus 3:431. 10.1186/2193-1801-3-431

Randi E, Lucchini V, Christensen M, Mucci N, Funk S, Dolf G, Loeschcke V (2001). Mitochondrial DNA Variability in Italian and East European Wolves: Detecting the Consequences of Small Population Size and Hybridization. Conserv Biol 14:464–473. 10.1046/j.1523-1739.2000.98280.x.

Randi E, Hulva P, Fabbri E, Galaverni M, Galov A, et al. (2014) Multilocus Detection of Wolf x Dog Hybridization in Italy, and Guidelines for Marker Selection. PLoS ONE 9(1): e86409. doi:10.1371/journal.pone.0086409

Ryabov L. (1973) Wolf-dog hybrids in the Voronezh oblast. Bull Mosk Soc Nat Biol 78(6): 25–39. (in Russian)

Salado I, Preick M, Lupiáñez-Corpas N, et al. (2022) Loss of Mitochondrial Genetic Diversity despite Population Growth: The Legacy of Past Wolf Population Declines. Genes 14:75. 10.3390/genes14010075.

Salvatori V, Donfrancesco V, Trouwborst A, et al. (2020). European agreements for nature conservation need to explicitly address wolf-dog hybridisation. Bioll Conserv 248. 108525. 10.1016/j.biocon.2020.108525

Saccone C, Attimonelli M, Sbisá E (1987). Structural elements highly preserved during the evolution of the D-loop-containing region in vertebrate mitochondrial DNA. J Mol Evol 26:205–11. 10.1007/BF02099853.

Sastre N, Vilà C, Salinas M. et al. (2011) Signatures of demographic bottlenecks in European wolf populations. Conserv Genet. 12:701–712. 10.1007/s10592-010-0177-6

Shibuya H, Collins BK, Huang TH-M, Johnson GS (1994) A polymorphic AGGAAT, tandem repeat in an intron of the canine von Willebrand factor gene. Animal Genet 25:122. 10.1111/j.1365-2052.1994.tb00094.x

Smeds L, Aspi J, Berglund J, Kojola I, Tirronen K, and Ellegren H (2021) Whole-genome analyses provide no evidence for dog introgression in Fennoscandian wolf populations. Evol Appl 14:721–734. 10.1111/eva.13151

Sundqvist AK, Ellegren H, Olivier M, Vilà C (2001) Y chromosome haplotyping in Scandinavian wolves (Canis lupus) based on microsatellite markers. Mol Ecol 10, 1959–1966. 10.1046/j.1365-294X.2001.01326.x

Suutarinen J, Kojola I (2017) Poaching regulates the legally hunted wolf population in Finland. Biol Conserv 215:11–18. 10.1016/j.biocon.2017.08.031

Talala MS, Bondarev AY, Zakharov ES, Politov DV (2020) Genetic differentiation of the wolf Canis lupus L. populations from Siberia at microsatellite loci. Russ J Genet 56(1):59–68. 10.1134/S1022795420010123

Tamura K, Stecher G, Kumar S (2021) MEGA 11: molecular evolutionary genetics analysis version 11. Mol Biol Evol 38(7):3022–3027. 10.1093/molbev/msab120.

Tikkunen M, Kojola I (2020) Does public information about wolf (Canis lupus) movements decrease wolf attacks on hunting dogs (C. familiaris)?. Nat Conserv 42:33–49. 10.3897/natureconservation.42.48314.

Tirronen KF (2008) Some particularities of wolf (Canis lupus) predation on domestic dogs (C. familiaris) in Karelia // The Herald of Game Manag 5(2):133–137. (in Russian)

Tirronen KF, Kuznetsova AS, Panchenko DV (2023) Population Genetic Structure of the Wolf (Canis lupus L.) in Eastern Fennoscandia under Conditions of Intensive Hunting Pressure Based on mtDNA Analysis. Biol Bull 50(5):1051–1063. 10.1134/S1062359023602240

Trouwborst A (2014) Exploring the Legal Status of Wolf-Dog Hybrids and Other Dubious Animals: International and EU Law and the Wildlife Conservation Problem of Hybridization with Domestic and Alien Species. Rev of Eur Compar Inter Environ Law 23. 10.1111/reel.12052

Yudin VG (2013) The wolf of the Far East of Russia. Dalnauka. Vladivostok (in Russian)

van Oosterhout C, Hutchinson WF, Wills DP, Shipley P (2004) Micro-checker: software for identifying and correcting genotyping errors in microsatellite data. Mol Ecol Notes 4:535–538. 10.1111/j.1471-8286.2004.00684.x

Vilà C, Amorim IR, Leonard JA, et al, (1999) Mitochondrial DNA phylogeography and population history of the gray wolf Canis lupus. Mol Ecol 8:2089–2103. 10.1046/j.1365-294x.1999.00825.x

Vilà C & Wayne R (2001) Hybridization between wolves and dogs. Conserv Biol 13:195–198. 10.1046/j.1523-1739.1999.97425.x

Vilaça S, Donaldson M, Benazzo A, Wheeldon T, Vizzari T, Bertorelle G, Patterson B, Kyle C (2023) Tracing Eastern Wolf Origins From Whole-Genome Data in Context of Extensive Hybridization. Mol Biol and Evol 40(4). 10.1093/molbev/msad055

